# Integration of transcriptional signatures from brain tissue and plasma extracellular vesicles of a preclinical tauopathy mouse model

**DOI:** 10.64898/2026.05.06.723062

**Authors:** Tanzima Tarannum Lucy, A. N. M. Mamun-Or-Rashid, Daniel C Lee, Iliya Lefterov, Radosveta Koldamova, Nicholas Francis Fitz

## Abstract

Tauopathies, including Alzheimer’s disease, involve progressive neurodegeneration and sustained neuroinflammation. We present a multi-compartment transcriptomic atlas of 9.6-month-old PS19 tauopathy mice compared with wild-type (WT) controls (n=8/group), profiling cortical mRNA, cortical non-coding RNA (ncRNA), and plasma small extracellular vesicle (pEV) ncRNA. In the PS19 cortex, mRNA sequencing identified 917 differentially expressed genes (DEGs), with microglial deconvolution revealing a robust transition toward disease-associated microglia (DAM) gene signature and downregulation of genes involved in oxidative phosphorylation and cholesterol biosynthesis relative to WT. Cortical ncRNA profiling identified 466 differentially expressed ncRNAs, primarily circular RNAs (circRNAs; n=331). In pEVs, 822 ncRNAs were differentially abundant, of which 657 circRNAs were identified in PS19 compared to WT mice. Cross-compartment integration demonstrated that pEV miRNA gene targets functionally mirrored genes involved in the brain’s inflammatory and metabolic failure. We identified a core shared signature of 33 ncRNAs, including miR-5114 (up in brain, down in pEV), circ_0008242 and circ_0002153 (up in brain and pEV), and circ_0007688 (down in brain and pEV) differentially enriched across both brain and periphery in PS19 compared to WT mice. These results demonstrate that the pEV non-coding landscape effectively tracks central tau-mediated changes in the brain transcriptional response. This study identifies circRNAs as the most numerically perturbed ncRNA class and provides a foundation for non-invasive biomarker development in tauopathy.

## Introduction

Alzheimer’s disease (AD) and related tauopathies affect more than 55 million people worldwide, with projections estimating 150 million cases by 2050 [1]. These conditions are defined by the intraneuronal accumulation of hyperphosphorylated tau forming neurofibrillary tangles (NFTs), deposition of amyloid-beta (Aβ) plaques, progressive synaptic loss, and sustained neuroinflammation. The revised 2024 Alzheimer’s Association diagnostic criteria now define AD biologically as a continuum—moving from initial molecular changes to overt clinical dementia—which highlights the urgent need for early, accessible biomarkers that can track disease progression across this spectrum [2–4].

Tauopathies represent a heterogeneous group of neurodegenerative disorders characterized by the pathological aggregation of the microtubule-associated protein tau (MAPT) into NFTs [5, 6]. Under physiological conditions, tau promotes microtubule assembly and axonal transport [7]. However, in diseases like AD and frontotemporal dementia (FTD), tau undergoes aberrant post-translational modifications, most notably hyperphosphorylation and acetylation—leading to its detachment from microtubules, loss of function, and subsequent neurotoxicity [6, 8].

The P301S transgenic mouse (PS19 line) is a widely used model of primary tauopathy that expresses human tau with the P301S MAPT mutation under the murine prion protein promoter, as it recapitulates the spatiotemporal progression of tauopathy, including early synaptic loss, microgliosis, and late-stage neuronal death in their hippocampus, neocortex and entorhinal cortex [9–11]. By 6 months of age, these mice develop NFTs, followed by progressive neuronal loss by 9–12 months and a pronounced neuroinflammatory profile characterized by microgliosis and astrogliosis [11–14]. Transcriptomic studies using bulk RNA-seq and Weighted Gene Co-expression Network Analysis (WGCNA) have shown that the gene expression in the cortex of PS19 mice overlaps with evolutionarily conserved disease modules identified in postmortem human FTD and AD brains, reinforcing the translational relevance of this model [15–17].

A growing body of research has established that the transcriptomic landscape of tauopathy extends beyond protein-coding mRNAs. Non-coding RNA (ncRNA) species—including microRNAs (miRNAs), circular RNAs (circRNAs), small nucleolar RNAs (snoRNAs), small nuclear RNAs (snRNAs), and transfer RNAs (tRNAs)—are emerging as significant post-transcriptional regulators that are extensively perturbed in AD and related dementias [18–20]. circRNAs are abundantly expressed in the brain and enriched at synapses; their covalently closed loop structure makes them uniquely resistant to exonucleases, suggesting they may serve as stable markers of neuronal state. Recent studies using m6A-seq and circRNA-seq in Drosophila tauopathy models and induced pluripotent stem cells (iPSCs)-derived neurons have demonstrated that tau pathology drives N6-methyladenosine (m6A) RNA methylation, which in turn regulates circRNA biogenesis and promotes neurodegeneration [21, 22].

Extracellular vesicles (EVs) are lipid bilayer-enclosed nanoparticles released by CNS cells that can traverse the blood-brain barrier bidirectionally [23–27]. These EVs carry functional cargo, including proteins and ncRNAs, which reflect the molecular state of their cell of origin. Furthermore, the discovery of a “molecular bridge” between the brain and periphery via EVs offers a unique opportunity for non-invasive monitoring [3, 23, 24]. Previous work by our group has highlighted the diagnostic potential of plasma EV non-coding RNA cargos, identifying snoRNAs as potential plasma EV biomarkers for AD and demonstrating that circulating EVs are essential mediators of neuroprotective signaling and cognitive resilience [28–31]. Human clinical studies using immuno-affinity enrichment to isolate neuron-derived EVs (targeting markers like L1CAM) from the plasma samples have shown that these vesicles carry tau species and miRNAs that can discriminate between AD and controls with high accuracy [32]. Furthermore, miRNA enrichment profiles in EVs have been shown to correlate with temporal cortical thickness on MRI in AD patients [32].

Despite these advances, several knowledge gaps persist. Most studies have focused on single compartments, and no study has simultaneously profiled the brain mRNA, brain ncRNA, and plasma EV ncRNA landscapes in the PS19 model. This hinders our understanding of how central molecular changes are reflected in peripheral biofluids. We hypothesize that tau pathology in the PS19 brain induces a coordinated ncRNA response that is packaged into EVs and detectable in the plasma, mirroring the central nervous system. By integrating transcriptomic data across central and peripheral compartments, we aimed to identify a convergent molecular signature with high fidelity for tracking the progression of tau-mediated neurodegeneration.

## Materials and Methods

### Animals

The PS19 mouse strain (B6;C3-Tg(Prnp MAPT*P301S)PS19Vle/J; (C57BL/6 × C3H)F1) was purchased from The Jackson Laboratory (Bar Harbor, ME, USA) and were used experimentally as heterozygous. Wild-type (WT) littermates were internally bred. All animal studies were approved by the University of Pittsburgh Institutional Animal Care and Use Committee (IACUC) and conducted in accordance with the guidelines of the Care and Use of Laboratory Animals. Mice were housed in a 12:12 h light–dark cycle with unrestricted access to food and water. Eight mice per group (equal number of male and female per group) were used for all experiments.

### Tissue and Plasma Processing

Mice were anesthetized with Avertin (1.25% tribromoethanol, 2.5% 2-methyl-2-butanol; 250 mg/kg, intraperitoneally). Blood was collected intracardially using an EDTA-treated syringe, followed by transcardial perfusion with chilled 20 mL of 0.1 M PBS (pH 7.4). Plasma was obtained by centrifugation at 13,000 × g for 5 min. For brain tissue RNA isolation, one hemisphere was dissected to use ∼10 mg cortex after removing the cerebellum, olfactory bulb, and subcortex. The other hemisphere was fixed in 4% phosphate-buffered paraformaldehyde at 4 °C for 48 h before being transferred to 30% sucrose for storage at 4 °C. Fixed hemibrains were embedded in O.C.T., sectioned coronally at 30 µm using a cryostat (Thermo Scientific), and stored in a glycol-based cryoprotectant at −20 °C until histological staining. All other samples were snap-frozen in dry ice before long-term storage at −80 °C.

### RNA Isolation

Brain mRNA and ncRNA were isolated from 10 mg of prefrontal cortex using the Norgen MicroRNA Isolation Kit (#21300, Canada). Tissue was dissociated in RLT buffer (#74106, Qiagen RNeasy Mini Kit, Germany) with 10% beta-mercaptoethanol, homogenized through a 25G needle, and passed through a QIAshredder (#79656, Qiagen, MD) prior to RNA isolation per manufacturer’s protocol. pEVs were isolated from 200 µL plasma using the Total Exosome Isolation Kit (#4484450, Invitrogen, Vilnius, Lithuania) following manufacturer’s protocol. EV RNA was extracted using the ExoQuick Exosome RNA Purification Column Kit (EQ808A, SBI, CA). RNA quality and quantity were assessed using the RNA Pico 6000 Kit for the Agilent 2100 Bioanalyzer (part ID: 5067-1513, Waldbronn, Germany).

### EV Characterization

EVs were analyzed by MRPS using the nCS2 instrument (Spectradyne, CA, USA) with C-400 microfluidic cartridges. Samples were diluted in 0.02 µm-filtered (Whatman Anotop 25/0.025, #6809-2102, Cytiva, Dassel, Germany) 1% Tween-20 in PBS and analyzed for a minimum of 30 runs with default settings. Western blotting was performed using EV lysates with cOmplete, EDTA-free Protease Inhibitor Cocktail (#11873580001) and Roche PhosSTOP (#4906837001) in 4–12% Bis-Tris WedgeWell gels (#NW04125BOX, Invitrogen, CA) transferred to nitrocellulose membranes (iblot 3 transfer stack, #IB33001, Invitrogen, Israel) using iblot3 transfer system (#IB31001, Invitrogen, Singapore). These membranes were washed and blocked in TBS Blocking buffer (#37579, Thermoscientific, IL), then probed with EV specific antibodies, anti-CD63 (#PA5-92370, Invitrogen, IL, 1:500 dilution o/n at 4°C), followed by goat anti-rabbit HRP (#A27036, Invitrogen, India, 1:10,000 dilution for 1 h at RT), and anti-CD81 (sc-7637, 1:200 o/n at 4°C), followed by goat anti-Mouse HRP (A28177, Invitrogen, India, 1:3,000 dilution for 1 h at RT). Also, we probed for endoplasmic reticulum (ER) specific Calnexin (ab22595, abcam, UK, 1:1000 dilution o/n at 4°C), followed by goat anti-rabbit HRP (A27036, 1:10,000 for 1 h at RT). Immunoreactive signals were visualized using enhanced chemiluminescence (TMA-6, Lumigen ECL Ultra, MI) with the Amersham Imager 600 (GE Lifescience).

### Library Preparation and Sequencing

mRNA libraries were generated using the SMART-Seq mRNA Kit (#634773, Takara, CA) combined with the Nextera XT DNA Library Preparation Kit (#FC-131-1096, Illumina, CA) and Illumina DNA/RNA UD Indexes Set A (#20091654, Illumina, CA). Small ncRNA libraries were prepared using the NEBNext Multiplex Small RNA Library Prep Set (#E7560S, NEB, MA). All libraries were sequenced on the NextSeq 2000 (Illumina, CA) generating paired-end reads at the Health Sciences Sequencing Core, University of Pittsburgh Children’s Hospital.

### Bioinformatics Analysis

Brain mRNA data were aligned to the mouse reference genome (mm10) using Rsubread (v2.16.1) [33]and annotated with org.Mm.eg.db (v3.19.1). Differential expression was analyzed with edgeR (v4.0.16) [34]. ncRNA libraries were aligned and quality-checked using STAR (v2.5.3a) with COMPSRA (v1.0.3) [35] and differential abundance determined by DESeq2 (v1.42.1) [36]. Genes and ncRNAs with *p* < 0.05 were considered differentially expressed. For ncRNA, GO enrichment analyses were performed using DAVID 2025 web-tool [37, 38] with a *p*-value cut-off of <0.05. For mRNA analyses, functional relationships among differentially expressed genes were assessed using enrichment analysis performed with Metascape (v3.5) [39]. Significantly enriched Gene Ontology (GO) terms were subsequently visualized as a network in Cytoscape (v3.10.4) [40], where each node represents an individual GO term and edges indicate shared gene membership. Functional clusters within the network were identified using the MCODE plugin [35]. To enhance network clarity and visual interpretability, edge bundling and manual layout adjustments were applied in Cytoscape (v3.x) [40]. In the final network representation, nodes are color–coded according to their assigned functional clusters, and node size is scaled to reflect enrichment significance (*p*–value). circRNA associated genes were predicted using circbase [41]. miRNA target predictions were performed using TargetScanMouse Release 8.0 [42].

To investigate cross-tissue relationships between brain and extracellular vesicle (EV) ncRNA profiles, a correlation network was constructed in R (v4.x) [43]. Expression values for brain- and EV-derived ncRNAs were treated as separate feature matrices and all pairwise Spearman correlations between brain–EV ncRNA pairs were computed using cor.test (1.0.7) [44] (two-sided, continuity-corrected). Edges were retained if they met the following criteria: raw *p* < 0.05, |ρ| ≥ 0.75. Nodes with fewer than 10 cross-tissue interactions in the initial filtered edge set (p < 0.05, |ρ| ≥ 0.75) were excluded; a small number of retained nodes may fall below this threshold in the final network due to the single-pass application of the degree filter. (For example: Node A had 10 brain interactions initially. Node B (one of its EV partners) had only 8 and was dropped. Now Node A has 9 but it already passed the filter.) Within-tissue pairs (brain–brain and EV–EV) were excluded from analysis. The resulting network was constructed using igraph (v1.x) and visualized using ggraph (v2.x), with node color encoding tissue of origin (brain: green; EV: blue), node size scaled to the number of interactions, and edge color and width reflecting correlation direction and magnitude, respectively. The network was additionally exported in GraphML format for interactive exploration in Cytoscape (v3.10.4) [40], where node shape and fill color were mapped to tissue source and edge width to |ρ|.

### Immunohistochemistry

To characterize changes in activated microglia in the vicinity of the infusion site, a series of six brain sections from each animal was immunostained with anti-IBA1 antibody. Six coronal sections per mouse (3 male, 3 female) were immunostained using Alexa Fluor 594–conjugated anti-IBA1 antibody (48934, Cell Signaling Technology, 1:500). Free-floating sections were washed in PBS, subjected to antigen retrieval (50 mM sodium citrate, 0.05% Tween-20, pH 9, 80°C, 30 min), blocked in 5% normal rabbit serum (0.2% Triton X-100/PBS, 1 h), and incubated with primary antibody overnight at 4°C followed by 1 h incubation at RT. Slides were mounted and imaged at 20× magnification on a Nikon Eclipse 90i microscope equipped with pco.panda 4.2M camera (Germany). The number of IBA1-positive microglia were quantified per field (four fields per section, six sections per mouse) using Nikon NIS Elements software.

### Statistical analysis

Statistical analyses and visualizations were performed using GraphPad Prism (v10.0.3) and a *p*-value of <0.05 was considered as significant.

## Results

We conducted a multi-compartment transcriptomic study using cortical brain tissue and plasma small extracellular vesicles (pEVs) from 8 PS19 and 8 wild-type (WT) mice (equal number of males and females, ∼9.6 months old). The study design integrated protein-coding mRNA and non-coding RNA (ncRNA) sequencing data from the brain cortex as it is already reported to show tauopathologies *i.e.,* NFTs, neuronal loss and brain atrophy, microgliosis and astrogliosis [2, 5–7, 11, 14, 45] and pEV ncRNA sequencing data to map the molecular landscape of tauopathy across central and peripheral compartments (Figure 1).

**Figure 1:**
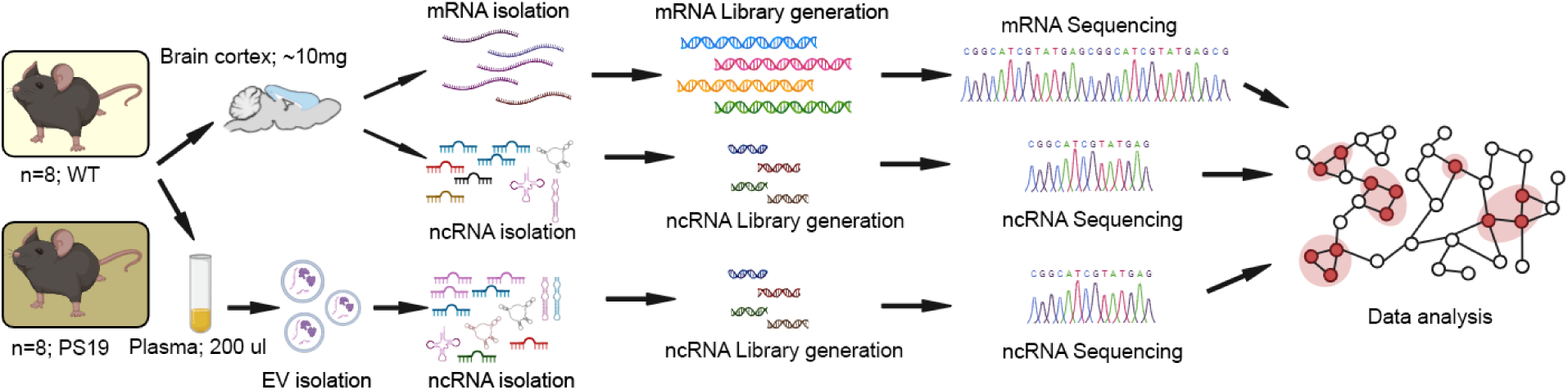
Schematic diagram of the study design. Brain cortex and plasma were collected from 8 WT and 8 PS19 mice (equal distribution of males and females, ∼9.6 months old). Cortical tissue (∼10 mg) was processed for parallel mRNA and small non-coding RNA (ncRNA) isolation. Plasma (∼200 μL) was used for EV isolation via precipitation-based methods, followed by ncRNA extraction. Following library generation and sequencing, data integration was performed to identify convergent molecular signatures across compartments. The figure was created using BioRender.com. (web-based; accessed on 2^nd^ April 2026).

### Cortical mRNA Differential Expression and Microglial Transcriptional Response in PS19 Mice

To investigate the primary transcriptomic alterations driving neurodegeneration in PS19, we first profiled the cortical mRNA landscape, hypothesizing that tau accumulation triggers cell-type specific inflammatory and metabolic shifts. Transcriptomic profiling of the prefrontal cortex identified 917 differentially expressed genes (DEGs) in PS19 mice compared to WT (568 upregulated and 349 downregulated in PS19 compared to WT; p < 0.05; Figure 2A). To determine the cellular contributions to this signature, we performed cell-type deconvolution based on our previous studies [46–52] which revealed that the highest number of microglia specific genes were the most significantly affected in PS19 mice compared to WT (Figure 2B). Further analysis of microglial specific DEGs showed a robust induction of disease-associated microglial (DAM) gene markers—including *Trem2*, *Gns*, *Man2b1*—and a corresponding downregulation of homeostatic gene markers such as *C1qa*, *C1qc*, *Tgfbr1* and *Csf1r* in PS19 mice compared to WT (Figure 2C). This homeostatic-to-DAM transcriptional transition reflects a conserved response to neurodegeneration documented in several models, such as 5xFAD and APP/PS1 mice, using single-cell RNA sequencing and immunohistochemistry to track microglial polarization [53–55].

**Figure 2:**
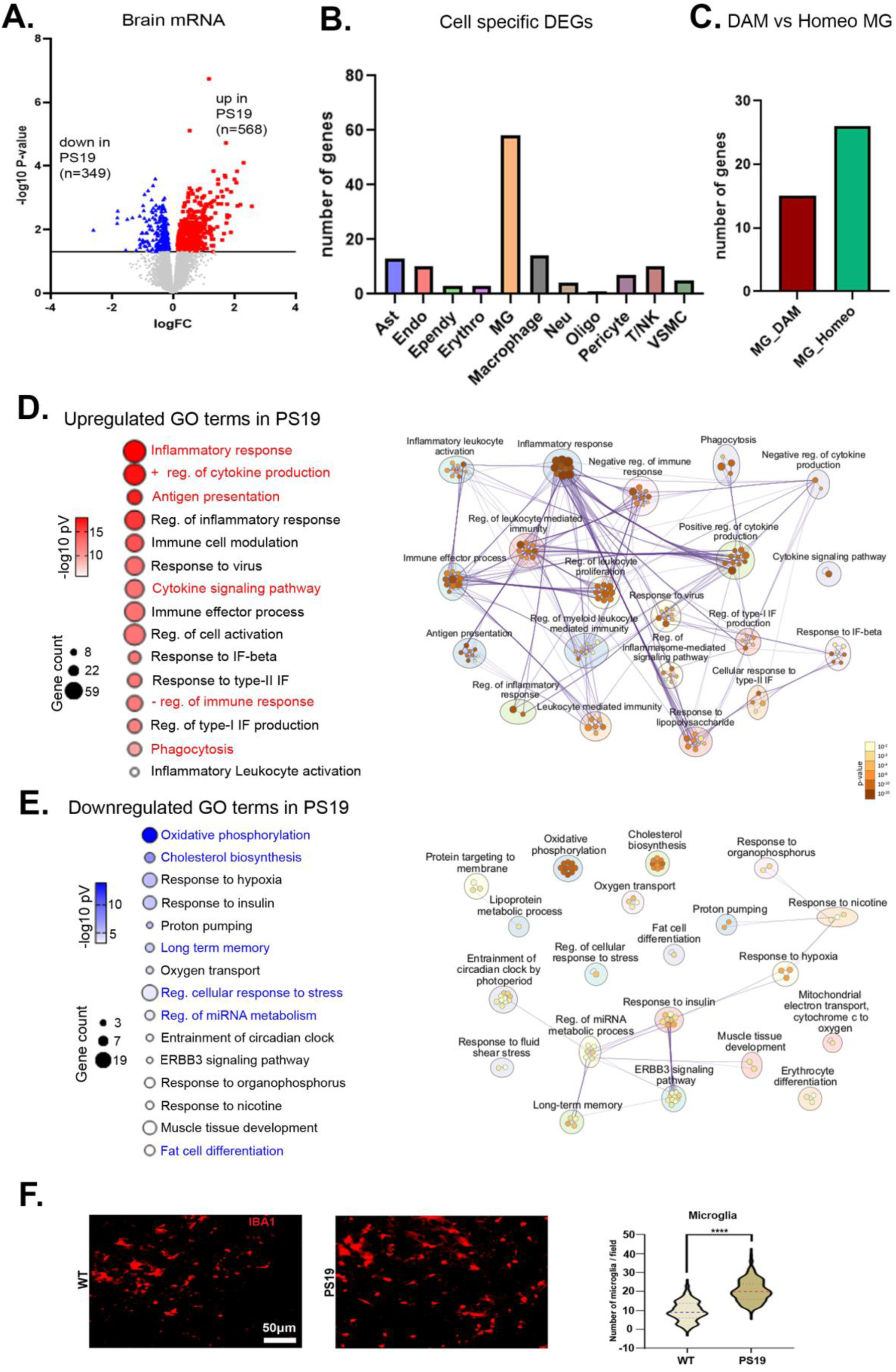
Cortical brain mRNA differential expression analysis shows upregulation of genes associated with inflammation and downregulation of genes associated with metabolism in PS19 mice compared to WT mice. Gene expression profiling was performed by RNA-seq followed by edgeR for identifying differentially expressed genes (DEGs) between cortical tissue from PS19 and WT mice. (A) Volcano plot showing differentially expressed cortical brain mRNAs in PS19 mice compared to WT (∼9.6 months old, n=8 per group); p < 0.05. Red, blue and gray denote significant upregulated (n=568) and downregulated (n=349) DEGs, and non-significant (n=13,400) genes, respectively. (B) Cell type-specific significant DEG distribution shows microglia specific genes are the most significantly affected in PS19 mice compared to WT. (C) Bar chart showing microglial state of DEGs; both DAM and Homeostatic genes were affected in PS19 mice compared to WT. (D, E) GO terms were analyzed using the Metascape [39] for the total upregulated and downregulated genes in PS19 mice compared to WT shown in (A). Bubble plots show GO Terms for upregulated and downregulated mRNAs respectively (left panels) in PS19 mice compared to WT. Bubble size represents gene count and color represents -log10(*p*-value). Enriched GO terms derived from DEGs were identified using Metascape and organized into functional modules. The resulting GO network was visualized in Cytoscape, where nodes represent enriched biological terms and node size reflects the number of associated input genes. Edges denote similarity between GO terms based on shared gene membership (kappa score > 0.3). Distinct node colors indicate separate functional clusters, highlighting modular organization within the gene set. (D, E-right panels). Immunohistochemistry data shows a significant increase in number of Ionized calcium-binding adaptor molecule 1 (IBA1)-positive microglia in the cortex of PS19 mice compared to WT. (F) Representative images of immunohistochemistry of IBA1 of the cortex of WT and PS19 brain sections (left) and violin plot showing the number of microglia per field. (∼6 months old, n=6 per group, equal number of male and female, four fields per section, six sections per mouse); statistical analysis was performed with unpaired t test. Plots are means ± SEM. ∗∗∗∗p < 0.0001.

Gene Ontology (GO) enrichment analysis indicated that total genes significantly upregulated in PS19 mice compared to WT were predominantly involved in ‘Inflammatory response’, ‘Antigen presentation’, ‘Cytokine signaling pathway’ and ‘phagocytosis’, and these upregulated pathways are closely associated with each other (Figure 2D). The upregulation of genes in these pathways is consistent with the NF-κB-driven microglial signature typically identified in tau-laden cortices of PS19 mice [56]. These modules share substantial overlaps with conserved neurodegenerative disease modules identified through WGCNA co-expression analysis in PS19 and rTg4510 mice and integrated with human postmortem FTD/AD data [15, 16]. Conversely, genes significantly downregulated in PS19 mice compared to WT were associated with ‘Oxidative phosphorylation’, ‘Cholesterol biosynthesis’, ‘Response to insulin’ and ‘Long-term memory’, unlike pathways associated with upregulated genes, only some of these pathways were closely associated with each other (Figure 2E). The suppression of oxidative phosphorylation represents a critical metabolic failure in PS19 compared to WT mice, likely driven by hyperphosphorylated tau-mediated impairment of mitochondrial transport as observed in various tauopathy models [4, 57, 58].

To validate the neuroinflammatory state, we performed immunohistochemistry (IHC) for ionized calcium-binding adapter molecule 1 (IBA1)-positive microglia in the cortex of younger mice (∼6 months old, n=6 per group). We observed a significant increase in the density of IBA1-positive microglia in PS19 cortex compared to WT (p < 0.0001; Figure 2F), providing biological confirmation of the DAM-associated transcriptomic profiles. Together, these data indicate that cortical tauopathy in the PS19 model is defined by a robust transition from homeostatic to disease-associated microglia alongside widespread downregulation of genes associated with metabolic functions compared to WT controls.

### Characterization of the Brain Non-Coding RNA Landscape in the PS19 Mouse Model

Given the substantial alterations in mRNA expression, we next analyzed the cortical ncRNA differential abundance to assess whether they act as key post-transcriptional regulators of the observed transcriptional changes and to what extent they reflect the mRNA expression patterns presented in Figure 2. We characterized the distribution and differential expression of five major ncRNA subtypes (circRNA, snoRNA, snRNA, miRNA, tRNA) in the cortex. Annotated counts across the classes revealed the broad diversity of the detected brain ncRNA landscape (Figure 3A). We identified 466 differentially expressed ncRNAs in PS19 mice compared to WT (219 downregulated and 247 upregulated in the cortex of PS19 mice compared to WT; Figure 3B). circRNAs constituted most of the differentially expressed fraction (n=331; 71% of total significant differentially expressed ncRNAs), highlighting them as the most numerically perturbed ncRNA class in the tauopathic brain of PS19 mice compared to WT (Figure 3C-D). We also observed significant perturbations in miRNAs (n=57; Figure 3G), tRNAs (n=30; Figure 3J), snoRNAs (n=46; Figure 3K), and snRNAs (n=2; Figure 3L) in PS19 mice compared to WT. This massive circRNA differential expression is a recently recognized feature of tauopathy, mechanistically linked to tau-induced N6-methyladenosine (m6A) RNA methylation-dependent biogenesis, a process validated using m6A-seq and circRNA-seq in Drosophila tauopathy models and human AD/FTD brain samples [21, 22].

**Figure 3:**
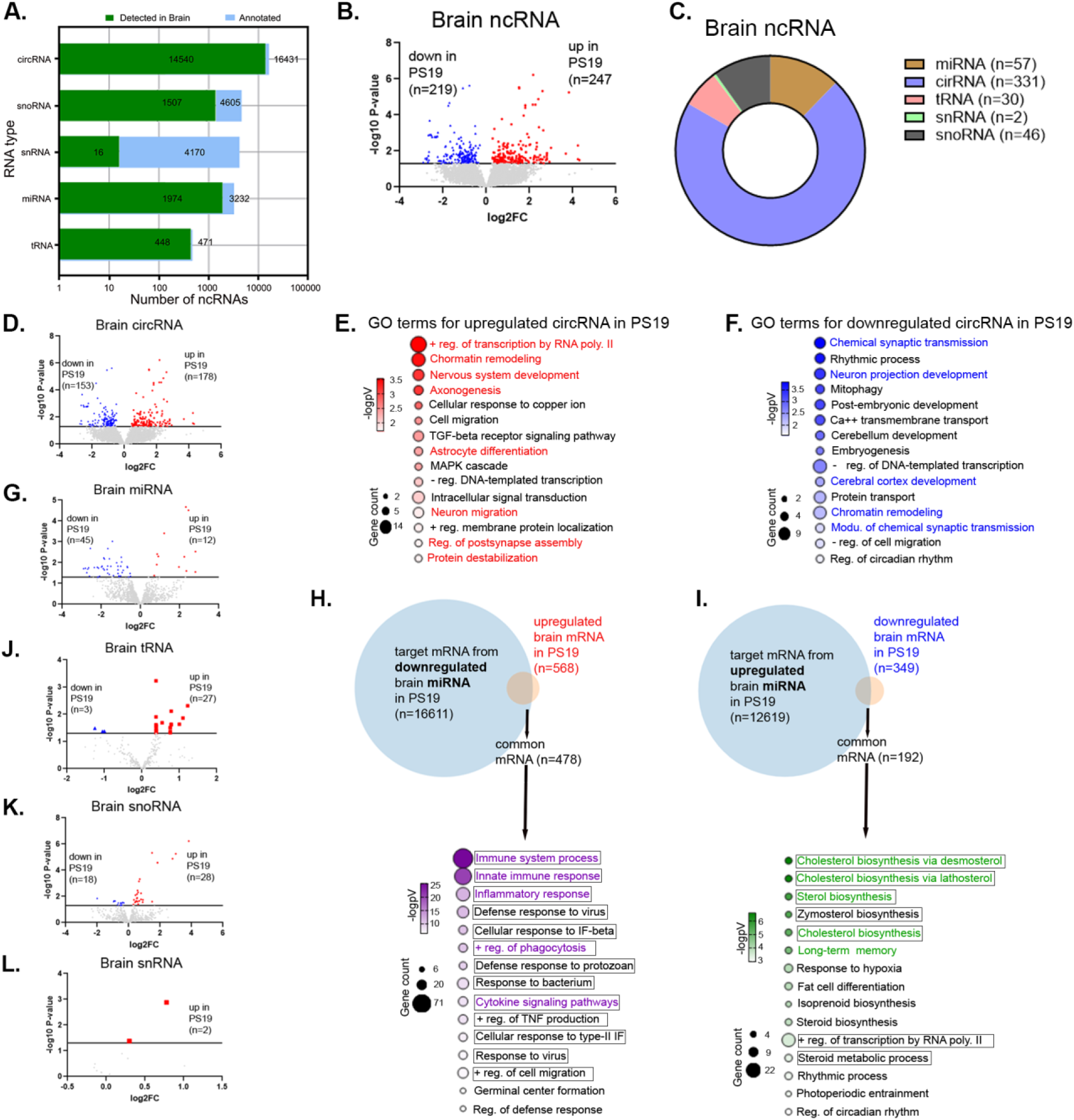
Differences in expression of cortical non-coding RNAs in PS19 mice and associated biological pathways of their host genes or predicted targets. (A) Bar graph shows the distribution of five ncRNA types (circRNA, snoRNA, snRNA, miRNA, and tRNA) from cortex based on annotated counts identified with COMPSRA. (B) Volcano plot of differentially expressed total brain ncRNAs comparing PS19 and WT mice (247 up and 219 down in PS19 mice), (D) circRNAs (178 up, 153 down), (G) miRNAs (12 up, 45 down), (J) tRNAs (27 up, 3 down), (K) snoRNAs (28 up, 18 down), and (L) snRNAs (2 up) in PS19 mice compared to WT. Red, blue and gray denote significant upregulated and downregulated differentially expressed ncRNAs, and non-significant ncRNAs in PS19 mice compared to WT, respectively. The results were considered significant with *p* < 0.05. (C) Donut chart summarizing significantly enriched [up + down; shown in (D), (G), (J), (K), (L)] ncRNA subtypes detected in brain tissue shown in (A). (E, F) DAVID was used to generate the bubble plots of GO enrichment analysis of the host genes (obtained from circBase) of upregulated (E) and downregulated (F) circRNAs. (H, I) Integration of significantly downregulated and upregulated brain miRNA targets (obtained using TargetScan mouse) with their corresponding brain mRNA data identified in Figure 2 (DEGs, up and down, respectively). Venn diagrams show common mRNAs between miRNA targets and their corresponding brain DEG lists. Bubble plots display GO terms for these common genes. Bubble size represents gene count; color represents -log10(*p*-value). Data shown upregulated or downregulated in PS19 mice compared to WT (∼9.6 months old, n=8 per group, equal number of male and female). non-coding RNAs (ncRNAs), microRNAs (miRNAs), circular RNAs (circRNAs), small nucleolar RNAs (snoRNAs), small nuclear RNAs (snRNAs), and transfer RNAs (tRNAs).

GO analysis of host genes associated with the 178 upregulated circRNAs in the PS19 cortex revealed enrichment in processes related to chromatin remodeling and transcriptional regulation, as well as neurodevelopmental programs including nervous system development, axonogenesis, and neuron migration (Figure 3E). These pathways are typically engaged during neuronal differentiation and synapse formation, suggesting reactivation of developmental and epigenetic programs that may reflect an attempt to maintain or reorganize neuronal identity in the context of ongoing synaptic stress. In contrast, host genes associated with the 153 downregulated circRNAs were predominantly enriched for processes directly linked to synaptic structure and function, including chemical synaptic transmission, neuron projection development, and cerebral cortex development, alongside chromatin remodeling (Figure 3F). The loss of circRNAs associated with these categories points more directly to impaired synaptic signaling and neuronal connectivity in the PS19 brain cortex. Taken together, although the upregulated and downregulated circRNA-associated gene sets initially appear to highlight distinct biological themes, both converge on pathways intimately linked to synaptic integrity and neuronal plasticity, consistent with progressive synaptic dysfunction in tauopathy. To evaluate miRNA-mediated regulatory effects, we integrated predicted mRNA targets of 45 downregulated miRNAs in the PS19 cortex (n = 16,611) with upregulated mRNAs (n = 568), identifying 478 shared genes associated with immune- and inflammation-related processes, including immune system response, inflammatory response, positive regulation of phagocytosis, and cytokine signaling pathways (Figure 3H). These pathways are hallmarks of genes related to microglial activation and innate immune signaling, indicating that loss of specific miRNAs may release repression on inflammatory gene networks in the PS19 cortex. Conversely, integration of mRNA targets of 12 upregulated miRNAs (n = 12,619) with downregulated mRNAs (n = 349) revealed 192 common genes enriched for metabolic and neuronal plasticity–related processes, including cholesterol and sterol biosynthesis, isoprenoid biosynthesis, long-term memory, and regulation of transcription by RNA polymerase II (Figure 3I). These pathways are essential for membrane homeostasis, synaptic function, and transcriptional control, suggesting that increased miRNA expression contributes to the suppression of genes related to core metabolic and cognitive-supporting programs in the PS19 cortex. Together, this integrated miRNA–mRNA regulatory landscape indicates a coordinated shift toward upregulation of genes related to neuroinflammatory signaling alongside repression of genes associated with metabolic and neuronal maintenance pathways, consistent with the transcriptomic alterations observed in Figures 2D and 2E. Thus, the cortical miRNA profile emerges as a key driver controlling genes involved in inflammatory activation and metabolic failure in the PS19 brain. In contrast to the circRNA expression signature, which primarily reflects genes associated with synaptic dysfunction and adaptive responses to synaptic loss, miRNA-mediated regulation appears to operate upstream by reshaping immune and metabolic gene networks that likely exacerbate neuronal vulnerability during tauopathy progression.

### Physical and Molecular Characterization of EVs Isolated from Plasma of PS19 and WT Mice

To assess whether central molecular perturbations are reflected in peripheral biofluids via ncRNAs, we performed physical and molecular characterization of pEVs, hypothesizing that their cargo mirrors the brain’s transcriptomic state. We isolated EVs from 200 μL of mouse plasma from both PS19 and WT mice using precipitation method described. Microfluidic Resistive Pulse Sensing (MRPS) analysis using the Spectradyne nCS2 confirmed a characteristic size distribution with a peak between 65–200 nm with no significant difference between WT and PS19 groups, aligning with MISEV2023 guidelines for defining small EV fractions [27] (Figure 4A). Total particle concentration was not different between groups with approximately 1 x 10^12^ particles/mL, and no significant differences were observed in distribution metrics D10, D50, or D90 (indicate the size below which 10%, the median particle size, and the size below which 90% of the particles fall, respectively) between WT and PS19 groups (p > 0.05; Figure 4B-C). This consistency aligns with clinical findings where pEV concentration remains stable while molecular cargo could change significantly in neurodegenerative diseases [59, 60].

**Figure 4:**
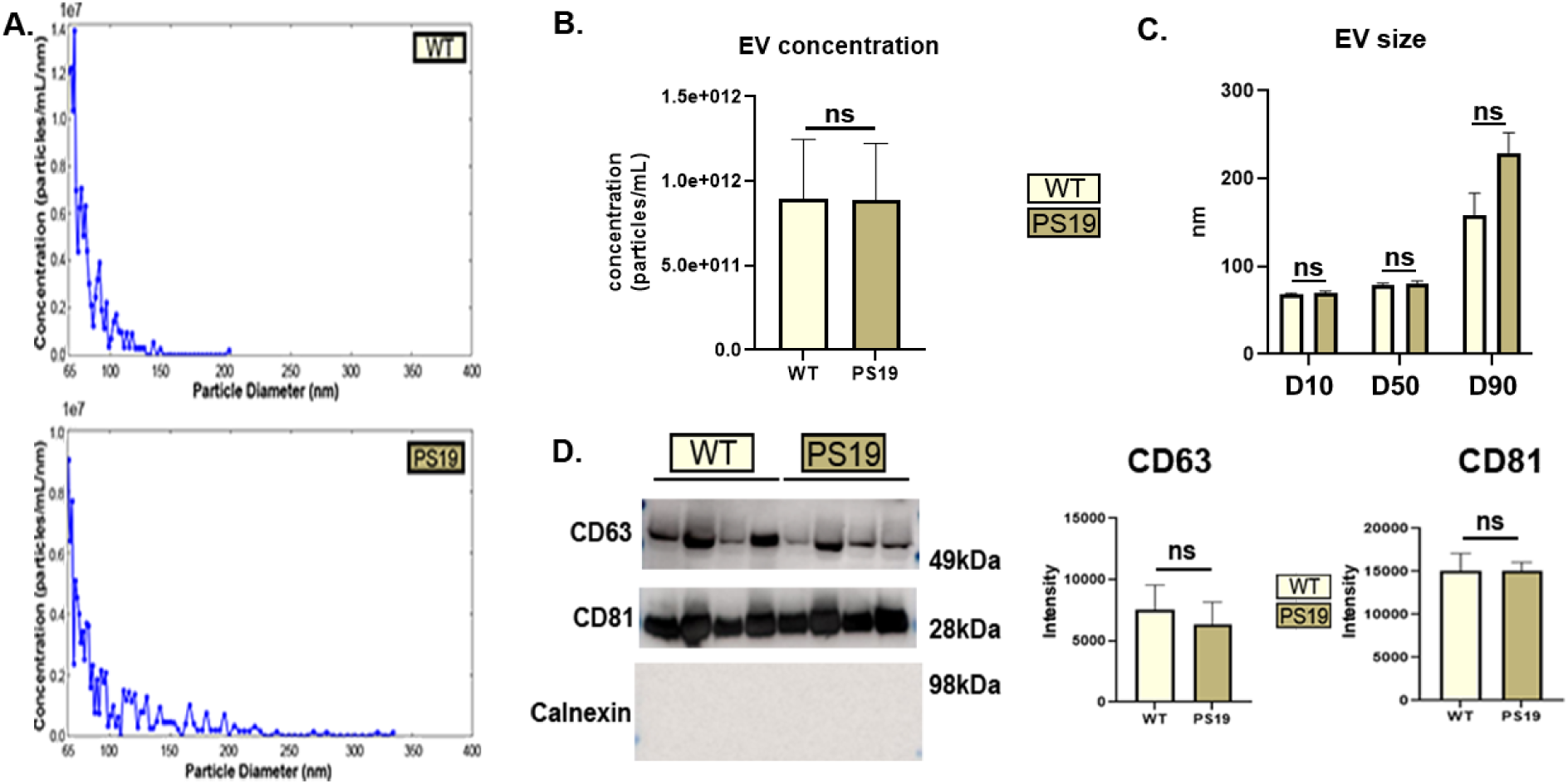
No difference in the characterization of EVs isolated from the plasma of PS19 and WT mice. (A) Representative Spectradyne particle analysis plots showing EV concentration and diameter in WT (top) and PS19 (bottom) mice. Quantification of Spectradyne particle analysis with bar graphs showing no significant (ns) difference in EV concentration (B) and EV size (C) between WT and PS19 mice. D10, D50 and D90 indicate the size below which 10%, the median particle size, and the size below which 90% of the particles fall, respectively. (D) Western blot of EV lysates showing EV tetraspanins markers, CD63 and CD81; and negative control marker, calnexin (left) and band intensity analysis for CD63 (center) and CD81 (right) showing no difference between the experimental groups. (∼9.7 months old, n=4, equal number of male and female per group). t-test was done for statistical analysis, ‘ns’ denotes ‘not significant’.

Western blot analysis (n=4 per group) confirmed the presence of canonical tetraspanin markers CD63 and CD81, with intensity quantification showing no significant difference in abundance between WT and PS19 pEV lysates, consistent with findings in other tau-transgenic mouse models [61, 62]. The absence of Calnexin confirmed the purity of the preparations and the exclusion of intracellular contaminants (Figure 4D). The stable physical characteristics of these pEVs across groups established a reliable baseline for the subsequent analysis of changes in their disease-specific molecular cargos.

### Detailed Profiling of ncRNA Cargo from EVs Isolated from Plasma of PS19 and WT mice

Following physical and molecular validation of pEVs, we next profiled their ncRNA cargo to assess whether circulating EVs reflect molecular alterations observed in the PS19 cortex when compared to WT. Annotation of pEV ncRNA subclasses detected total 15,157 ncRNAs (Figure 5A) and among them 822 were differentially enriched ncRNAs in PS19 mice compared to WT (581 upregulated and 241 downregulated; Figure 5B). As in the cortex, circRNAs represented the dominant differentially abundant species (n = 657; ∼80%), with a strong bias toward upregulation (518 circRNAs) in PS19 pEVs compared to WT (Figure 5C-D). Significant alterations were also observed in other ncRNA classes, including miRNAs (10 upregulated, 28 downregulated; Figure 5G), snoRNAs (6 upregulated, 74 downregulated; Figure 5J), and tRNAs (47 upregulated; Figure 5K), indicating broad but class-specific remodeling of the pEV ncRNA landscape in PS19 mice compared to WT.

**Figure 5:**
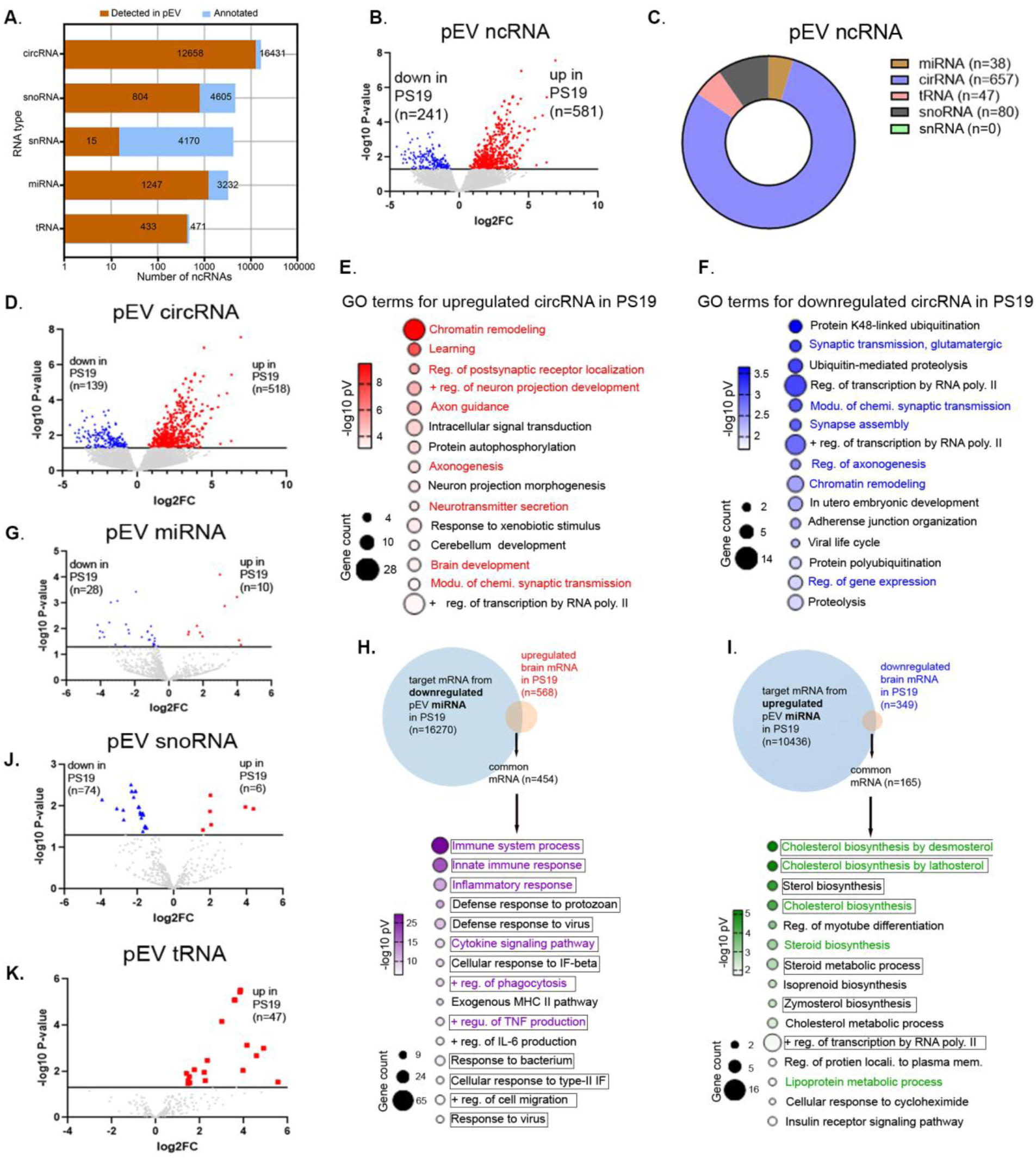
Differential Abundance and Functional Analysis of ncRNAs cargos from plasma EVs of PS19 and WT mice. (A) Bar graph shows the distribution of five ncRNA types (circRNA, snoRNA, snRNA, miRNA, and tRNA) in pEVs, based on annotated counts by COMPSRA. (B) Volcano plot of all differentially enriched ncRNA in pEVs of PS19 mice compared to WT mice (581 up, 241 down in pEVs of PS19 mice), (D) circRNA (518 up, 139 down), (G) miRNAs (10 up, 28 down), (J) snoRNAs (6 up, 74 down), and (K) tRNAs (47 up). Red, blue and gray denote significant upregulated and downregulated differentially enriched ncRNAs, and non-significant ncRNAs when comparing PS19 to WT groups, respectively. The results were considered significant when the *p* < 0.05. (C) Donut chart summarizing significantly enriched ncRNA subtypes [up + down; shown in (D), (G), (J), (K)] in pEV shown in (B). (E, F) DAVID was used to generate the bubble plots of GO enrichment analysis of the host genes (obtained from circBase) of upregulated (E) and downregulated (F) circRNAs. (H, I) Integration of significantly downregulated and upregulated pEV miRNA targets (obtained using TargetScan mouse) with their corresponding brain mRNA data from Figure 2 (DEGs, up and downregulated, respectively). Venn diagrams show common mRNAs between miRNA targets and their corresponding brain DEG lists. Bubble plots display GO terms for these common targets. Bubble size represents gene count; color represents - log10(*p*-value). Data shown upregulated or downregulated in PS19 mice compared to WT (∼9.6 months old, n=8 per group, equal number of male and female). non-coding RNAs (ncRNAs), microRNAs (miRNAs), circular RNAs (circRNAs), small nucleolar RNAs (snoRNAs), small nuclear RNAs (snRNAs), and transfer RNAs (tRNAs).

Functional enrichment analysis of host genes associated with upregulated circRNAs in PS19 pEVs highlighted pathways related to chromatin remodeling and transcriptional regulation, as well as neuronal and synaptic processes including learning, axon guidance, axonogenesis, postsynaptic receptor localization, and neuron projection development (Figure 5E). These terms are central to synaptic organization and plasticity, suggesting that circRNA cargo within pEVs captures neuronal responses to synaptic stress. Notably, this signature closely mirrors the enrichment profile of upregulated circRNAs in the PS19 cortex (Figure 3E), supporting the idea that circulating pEV circRNAs reflect central synaptic remodeling and degeneration-associated responses in tauopathy. GO analysis of host genes associated with downregulated circRNAs in PS19 pEVs revealed enrichment for pathways involved in synaptic organization and maintenance, including synaptic transmission, synapse assembly, and regulation of axonogenesis, alongside chromatin remodeling (Figure 5F). The coordinated loss of circRNAs linked to these processes is consistent with impaired synaptic integrity and mirrors the functional signature observed for downregulated circRNAs in the PS19 cortex compared to WT (Figure 3F), further supporting the convergence between circulating and brain-derived circRNA profiles.

Notably, downregulated circRNAs host genes were associated with protein K48-linked ubiquitination, highlighting disruption of genes important in the proteostatic machinery of PS19 mice. K48-linked ubiquitin chains serve as the canonical signal for proteasomal degradation, a pathway that is perturbed by toxic tau species, leading to impaired protein clearance. This finding aligns with evidence from Drosophila tauopathy models and Alzheimer’s disease postmortem brain tissue, where tau-associated proteasome dysfunction contributes to synaptic and neuronal degeneration [63–66]. Together, these data suggest that pEV circRNAs capture not only synaptic decline but also proteostasis defects that are central to tau-mediated neurodegeneration.

Integration of pEV miRNA gene targets with cortical mRNA expression revealed a strong functional concordance between circulating pEV associated ncRNAs and brain mRNA transcriptome. Predicted mRNA targets of 28 downregulated miRNAs from PS19 pEVs compared to WT (n = 16,270) overlapped with 454 upregulated mRNAs in the PS19 cortex and were enriched for genes associated with immune-related pathways, including immune system response, inflammatory response, regulation of phagocytosis, and cytokine signaling (Figure 5H). This pathway enrichment closely mirrors the regulatory signature observed for downregulated cortical miRNAs intersecting with upregulated mRNAs in PS19 mice (Figure 3H). Conversely, predicted mRNA targets of 10 upregulated miRNAs from PS19 pEVs (n = 10,436) intersected with 165 downregulated cortical mRNAs, revealing enrichment for genes associated with metabolic and transcriptional pathways such as cholesterol and steroid biosynthesis, isoprenoid biosynthesis, and regulation of transcription by RNA polymerase II (Figure 5I). These functional categories directly recapitulate the GO terms identified for upregulated cortical miRNAs and downregulated mRNAs in the PS19 cortex (Figure 3I). Collectively, these data indicate that the ncRNA cargo of circulating pEVs—particularly miRNAs—serves as a peripheral marker of the metabolic suppression and inflammatory activation occurring in the tauopathic brain[67–73]. Together with the circRNA findings, these results demonstrate that pEV ncRNA profiles encode a functional signature that closely reflects brain inflammatory and metabolic landscape of PS19 mice compared to WT mice, supporting their potential utility as biomarkers of tau-driven neurodegeneration.

### Integration of Functional Analysis of ncRNAs in Brain and pEVs from PS19 Compared to WT Mice

Network-based analysis of significant differentially changed ncRNAs (excluding the uncharacterized GM subtypes) revealed a highly coordinated relationship between ncRNAs from cortical brain tissue (green nodes) and circulating pEVs (blue nodes) across groups (Figure 6A). The resulting correlation network was dominated by negative correlations (blue edges), indicating that changes in cortical ncRNA expression levels were inversely related in pEV ncRNA cargos, while positive correlations (red edges) were less frequent. Within this network, miR–203 and miR–487b emerged as central hub nodes, positioning them as key mediators of brain–periphery communication. The strong negative correlation of these miRNAs between cortex and pEVs suggests that their peripheral abundance may serve as a surrogate biomarker of ncRNAs that regulate genes important in the brain inflammatory state, highlighting specific ncRNA clusters as potential non-invasive ‘liquid biopsy’ candidates for neurodegenerative conditions.

**Figure 6:**
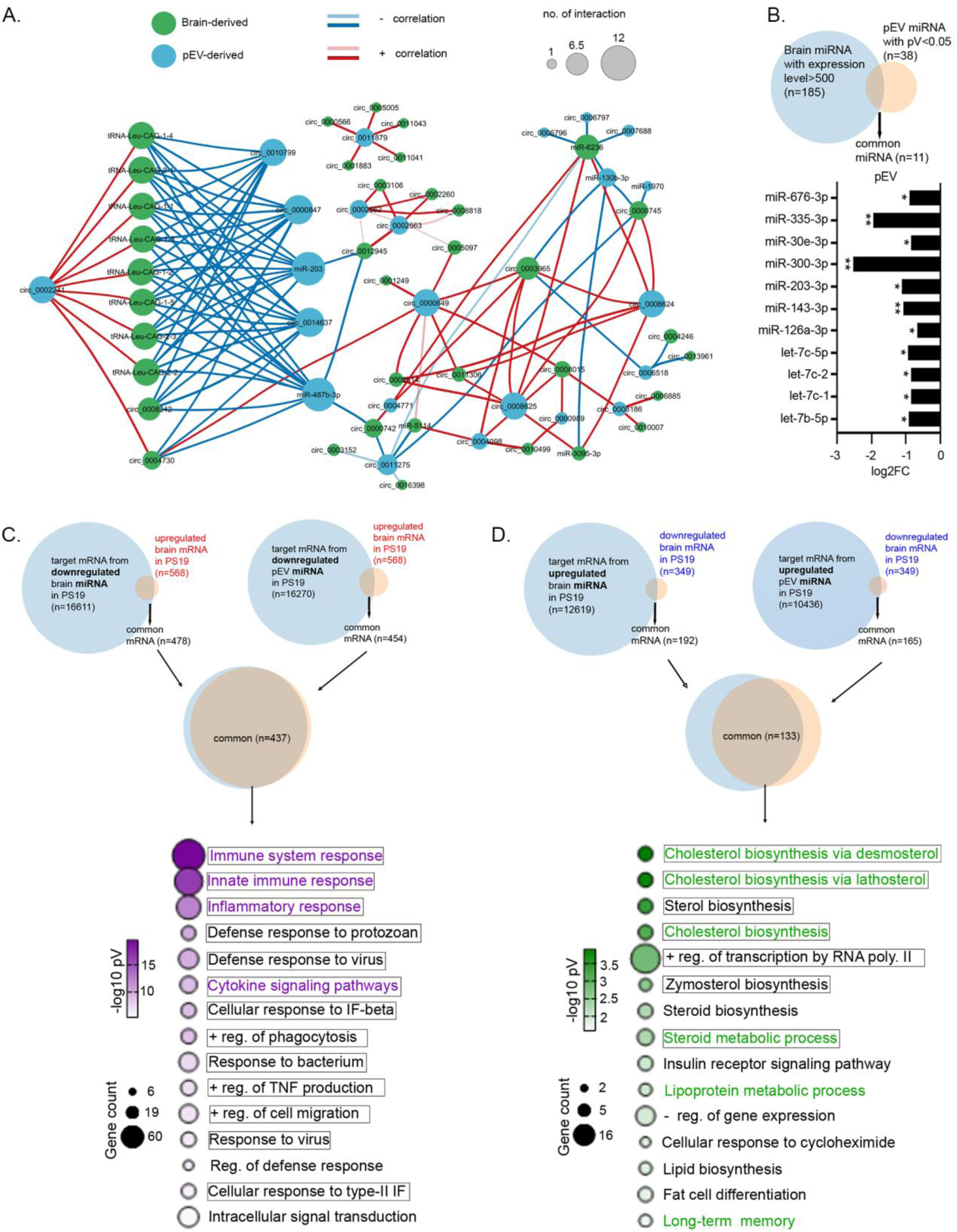
Comparative pathway analysis and integration of brain and pEV ncRNA signatures associated with tau pathology. (A) The correlation network diagram visualizes the complex interaction between significant differentially enriched ncRNAs in cortical brain tissue and pEVs of PS19 versus WT mice. A total of 38,140 significant correlations were observed among them 24,237 positive correlations and 13,903 negative correlations were observed; within-compartment pairs were excluded. Correlation pairs were then filtered by *p*-value, correlation strength, and minimum node degree to retain the most robust cross-tissue associations. Edges represent significant pairwise Spearman correlations (*p* < 0.05, |ρ| ≥ 0.75) between brain and pEV ncRNA pairs. The nodes are color-coded by compartment: green circles represent brain-derived ncRNAs, while blue circles represent pEV-derived ncRNAs. The size of each node reflects its number of interactions in the network, with larger circles indicating molecules that share numerous significant correlations with other transcripts across both compartments. Connections signify statistically significant correlations between pairs of ncRNAs. Red lines indicate a positive correlation, where higher expressions in brain correspond to higher abundance in pEVs. Blue lines indicate a negative correlation, where higher expressions in brain correspond to lower abundance in pEVs. The thickness of the lines represents the strength of the correlation, light thin lines indicate lower correlation, and dark thick lines indicate higher correlation. Key transcripts, including specific circRNAs, microRNAs, and tRNA fragments. (B) Venn diagram showing the overlap of brain miRNA with expression level > 500, considered brain enriched miRNAs, and significant differentially abundant (p <0.05) miRNA species in pEVs, identifying 11 common miRNAs. Bottom panel shows enrichment profile of those 11 shared miRNAs in pEVs of PS19 compared to WT. ∗*p* < 0.05, ∗∗*p* < 0.01. (C, D) Cross-compartment integration analysis. Venn diagrams illustrate the common miRNA gene targets (identified via TargetScan mouse) that are shared between significant differentially expressed brain miRNAs and differentially abundant pEV miRNAs. These shared gene targets were then overlapped with the cortical brain mRNA dataset. (C) Integration with upregulated brain mRNAs highlights shared genes associated with regulatory control of immune system response and cytokine-mediated signaling. (D) Integration of miRNA targets with downregulated brain mRNAs reveals highly convergent enrichment of genes associated with cholesterol and sterol biosynthetic processes, as shown in the accompanying bubble plot. Data shown upregulated or downregulated in PS19 mice compared to WT (∼9.6 months old, n=8 per group, equal number of male and female). non-coding RNAs (ncRNAs), microRNAs (miRNAs), circular RNAs (circRNAs), and transfer RNAs (tRNAs).

To further assess brain origins of pEV miRNA cargos, we compared highly expressed cortical miRNAs (average expression > 500 CPM; n = 185) with 38 significant differentially abundant pEV miRNAs, identifying 11 miRNAs that are brain enriched and differentially abundant in pEVs of PS19 mice compared to WT (Figure 6B). Notably, this group included miR–203–3p, reinforcing its role as a brain-derived mediator of peripheral signaling, consistent with its central hub node in Figure 6A.

Functional annotation of 437 shared target mRNAs derived from downregulated miRNAs in the cortex (n = 16,611) and pEVs (n = 16,270), intersected with 568 upregulated cortical mRNAs, revealed consistent enrichment of genes associated with immune-related processes, including immune system response, inflammatory response, cytokine signaling pathways, and regulation of phagocytosis across both compartments (Figure 6C). In contrast, 133 shared target mRNAs associated with upregulated miRNAs in the cortex (n = 12,619) and pEVs (n = 10,436), intersecting with 349 downregulated cortical mRNAs, were enriched with genes associated with pathways related to cholesterol and sterol biosynthesis, positive regulation of transcription by RNA polymerase II, and long-term memory (Figure 6D). Together, these analyses demonstrate that ncRNA abundance patterns in circulating pEVs robustly recapitulate the transcriptional responses associated with inflammatory activation and metabolic repression observed in the PS19 brain compared to WT. This convergence across compartments further supports the concept that pEV ncRNA profiles—particularly miRNA networks—serve as peripheral biomarkers of brain pathological processes during tauopathy.

### Shared ncRNA Signatures Between the Brain and pEV of PS19 mice Compared to WT

Finally, we integrated transcriptomic datasets across cortical tissue and circulating pEVs to identify convergent molecular signatures and assess the capacity of pEV-derived ncRNAs to reflect central tauopathy-associated changes. This analysis revealed a discrete shared signature of 33 ncRNAs that showed significant differential expression in brain and differential abundance in pEVs of PS19 mice compared with WT (Figure 7). This overlapping signature was overwhelmingly dominated by circRNAs (32 of 33), most of which exhibited concordant differentially enrichment in both brain and pEVs. Notably, circRNAs such as circ_0008242 and circ_0002153 were consistently upregulated in both brain and pEVs, whereas circ_0007688 was significantly reduced in both compartments (Figure 7–1i–ii).

**Figure 7:**
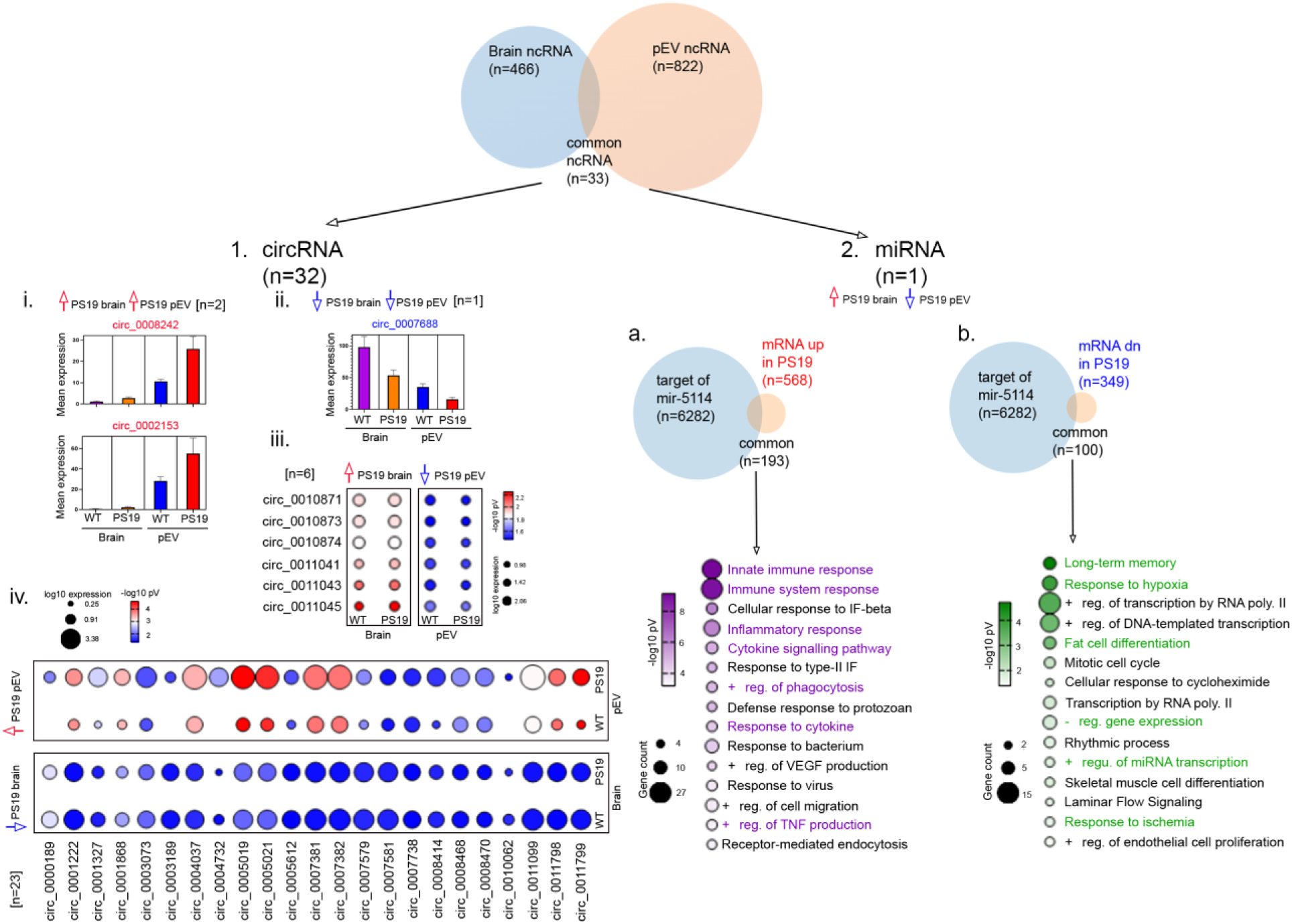
Common differentially enriched ncRNA species in both brain and pEVs of PS19 mice. (Top) Venn diagram illustrates 33 ncRNAs are both significant differentially expressed in the cortex and differentially abundant in pEVs of PS19 mice compared to WT (p < 0.05). (1) Detailed breakdown of the 32 common significantly altered circRNAs from brain and pEVs of PS19 mice compared to WT. (I, II) Bar graphs showing the mean expression of circRNAs that are significantly upregulated (I) or downregulated (II) in both the brain and pEVs of PS19 mice compared to WT, respectively. (III, IV) Comparative bubble plots for the remaining shared circRNAs, where bubble size reflects mean expression and color reflects the log2 fold-change (log2FC) directionality. Panel III shows 6 circRNAs that were significantly upregulated in the cortex but downregulated in pEVs of PS19 mice compared to WT. Panel IV shows 23 circRNA that were downregulated in cortex but upregulated in pEVs of PS19 mice compared to WT. (2) Functional mapping of miR-5114, the only shared differentially enriched miRNA in both brain and pEVs of PS19 mice. (a, b) Venn diagrams showing the overlap of predicted miR-5114 gene targets with cortical mRNAs that are upregulated (a) or downregulated (b) in PS19 mice compared to WT. GO term bubble plots reveal that miR-5114 targets are genes involved in innate immune responses and long-term memory/hypoxia response pathways, suggesting miR-5114 acts as a key molecular bridge between central pathology and peripheral EV cargo. Data shown upregulated or downregulated in PS19 mice compared to WT (∼9.6 months old, n=8 per group, equal number of male and female). non-coding RNAs (ncRNAs), microRNAs (miRNAs), and circular RNAs (circRNAs).

In addition to concordant regulation, several circRNAs displayed inverse expression-enrichment patterns between brain and pEVs. Specifically, six circRNAs were upregulated in the brain but downregulated in pEVs (Figure 7–1iii), while twenty–three circRNAs were downregulated in the brain yet upregulated in pEVs of PS19 mice compared to WT (Figure 7–1iv), consistent with selective ncRNA export into the peripheral circulation during disease progression.

Among miRNAs, miR–5114 emerged as the sole differentially abundant species shared between brain and pEVs, exhibiting increased levels in the PS19 cortex and reduced abundance in pEVs isolated from PS19 mice relative to WT (Figure 7–2). Comparison demonstrated that 193 mRNAs were common between predicted miR–5114 gene targets (n = 6,282) and upregulated cortical mRNAs revealed association of genes important in innate immune and inflammatory pathways (Figure 7–2a). In contrast, analysis of 100 shared mRNAs intersecting with downregulated cortical mRNAs identified genes related to neuronal function and adaptation, including long–term memory, response to hypoxia, and positive regulation of transcription by RNA polymerase II (Figure 7–2b). Together, these findings identify miR–5114 as miRNA altered in both brain and pEVs of PS19 mice compared to WT and has common gene targets with cortical DEGs associated with inflammatory activation with repression of neuronal maintenance processes. More broadly, the convergence of circRNA– and miRNA–based signatures across brain and pEVs highlights a compact, disease–relevant ncRNA network with strong potential as a non–invasive biomarker axis for tauopathy.

## Discussion

The simultaneous profiling of cortical mRNA, ncRNA, and pEV ncRNA in PS19 mice was used to test the hypothesis that the central molecular landscape of tauopathy is reflected in peripheral biofluids through EV-mediated transport. Our findings reveal that tau pathology in the PS19 cortex is associated with a pronounced transcriptional response associated with neuroinflammation dominated by genes associated with microglial activation and DAM gene induction. Furthermore, the transcriptional response was associated with a metabolic alteration characterized by suppression of oxidative phosphorylation and cholesterol biosynthesis, and a remarkably broad ncRNA differentially enriched landscape in which circRNAs emerge as the most perturbed class in both brain tissue and circulating pEVs. These results suggest a conserved neurodegenerative transcriptional response and the potential of ncRNAs as both mechanistic drivers of gene expression and peripheral biomarkers of tauopathy.

### Conservation of Neuroinflammatory and Metabolic Transcriptomic Signature Across Neurodegenerative Models and Human Disease

The dominant transcriptomic features identified in the PS19 cortex—specifically microglial DAM induction, inflammatory pathway upregulation, and metabolic gene suppression in PS19 compared to WT mice—are robustly conserved across various models of neurodegeneration and human disease. The enrichment of upregulated brain DEGs in antigen processing and presentation, myeloid leukocyte activation, and positive regulation of phagocytosis is also consistent with tau fibril–induced microglial activation via TLR2/MyD88/NF-κB signaling [45], and supports the concept that tau aggregates are recognized as damage-associated molecular patterns (DAMPs) by innate immune receptors. While DAM activation is considered partially protective—enhancing phagocytic clearance of toxic tau species—chronic activation of these pathways drives tau spreading through small EV release of seeding-competent tau fragments [74–76], creating a vicious cycle of inflammation and progressive tauopathy.

Santos et al. used WGCNA co-expression analysis in PS19, rTg4510, and GRN-haplo-insufficient mice to identify co-expression modules that were highly preserved in human postmortem FTD and AD brains, demonstrating conservation of the microglial/inflammatory and neuronal/synaptic gene modules [15, 16, 77]. We also observed that the PS19 cortex represents an evolutionarily conserved response to neurodegeneration. The downregulation of genes associated with oxidative phosphorylation in PS19 mice compared to WT reflects a metabolic failure common to human AD, often attributed to hyperphosphorylated tau-mediated disruption of the electron transport chain and mitochondrial transport impairment as shown in transgenic mouse models using proteomics and functional assays [45, 53–55, 57, 78]. Furthermore, the suppression of cholesterol biosynthesis in PS19 cortex compared to WT is significant, as cholesterol homeostasis is essential for synaptic integrity and its disruption is linked to accelerated tau phosphorylation in human-derived iPSC models [79–82]. The co-suppression of long-term memory–related gene programs alongside metabolic failure suggests that bioenergetic insufficiency may underlie early cognitive deficits in PS19 mice, a hypothesis supported by spatial transcriptomic data showing early downregulation of genes associated with ATP metabolic process in vulnerable hippocampal subfields [83]. These findings confirm that the PS19 model recapitulates the fundamental transcriptomic shifts observed in human tauopathy, providing a valid framework for biomarker discovery.

### CircRNA Differential Expression as a Significant Feature of Tauopathy ncRNA Biology

The increased differential enrichment of circRNAs in both the brain cortex (71%) and pEVs (80%) of PS19 mice compared to WT is the most striking finding of this study and confirms the importance of this subtype in the tauopathy landscape. circRNAs are stabilized by their covalently closed structure and are enriched at synapses where they regulate neuronal plasticity; their perturbation in the PS19 brain suggests a fundamental shift in synaptic regulatory architecture [84–90]. A mechanistic link between tau pathology and circRNA biogenesis was established by Atrian et al. [21], who used m6A-seq and circRNA-seq in Drosophila tauopathy models and iPSC- derived neurons to demonstrate that tau-induced N6-methyladenosine (m6A) RNA methylation drives circRNA biogenesis, leading to neurotoxic circRNA elevation. Our data extend these findings to the PS19 mammalian model and in both brain and pEVS, showing that hundreds of circRNAs are altered in PS19 mice compared to WT, with a notable preference toward downregulation in pEVs (518 species downregulated in PS19 pEVs compared to WT). This m6A-circRNA axis may explain why circRNA differential expression is so extensive in our PS19 dataset: widespread tau-induced m6A modifications could trigger global alterations in back-splicing rates across hundreds of loci [21, 22]. circRNAs can sponge miRNAs and thereby inhibit them from downregulating the translation of their target mRNAs. Upregulated brain circRNAs may sponge miRNAs that would eventually suppress synaptic transmission genes, while downregulated circRNAs may release suppression of chromatin remodeling and neuronal structural genes—resulting in a bidirectional disruption of neural circuitry maintenance. The enrichment of downregulated pEV circRNA host genes associated with ‘Protein K48-linked ubiquitination’ is of biological significance, as the proteasomal degradation pathway is a primary target of tau-mediated toxicity across models [19, 91–93]. Selective loading of circRNAs into EVs is a regulated process involving EV biogenesis machinery and altered circRNA packaging may also reflect tau-driven changes in multivesicular body biology or altered expression in source cells (neurons, microglia, endothelial cells) contributing to the circulating pEV pool. Studies profiling circRNAs in plasma of AD patients have identified differential circRNA signatures that could discriminate AD from MCI and healthy controls [20], supporting the translational relevance of our findings. The GO pathway convergence between brain and pEV circRNA host gene networks, particularly for synaptic transmission and neurodevelopmental pathways—further support the conclusion that pEV circRNA cargo reflects genuine brain transcriptional changes rather than peripheral systemic responses.

### miRNA Regulatory Networks Linking Brain and Plasma EVs in PS19 mice

Our miRNA integration analyses reveal a coherent picture in which both brain-derived and pEV-derived miRNAs have target genes which converge on the suppression of genes associated with cholesterol/sterol biosynthesis and amplification of immune and inflammatory responses in PS19 mice compared to WT. This convergence suggests that miRNAs loaded into plasma EVs are not a random sampling of brain miRNA expression but may include species specifically exported via EV biogenesis pathways under pathological conditions.

The identification of miR-126a-3p, miR-335-3p, and miR-300-3p, among the most differentially abundant miRNAs in pEVs that are also highly expressed in the brain, are of particular interest. miR-126-3p is a well-established regulator of vascular integrity and angiogenesis, predominantly expressed by endothelial cells, and has been reported to be significantly downregulated in plasma-derived EVs of AD patients [67–73]. Its shared differentially enrichment across brain tissue and circulating pEVs in PS19 mice raises the possibility that endothelial cell–derived EVs carrying miR-126 reflect tau-associated vascular compromise, consistent with evidence of blood–brain barrier disruption in PS19 mice and in human tauopathy. miR-335-3p has been shown to regulate cellular stress responses and has been identified in panels of AD-responsive peripheral miRNAs with reported roles in regulating synaptic plasticity genes relevant to memory consolidation [94]. The co-regulation of genes important in cholesterol biosynthetic pathway by both brain miRNAs and pEV miRNAs—where upregulated miRNAs target cholesterol biosynthesis genes whose mRNAs are simultaneously downregulated in the brain transcriptome—suggests an integrated multi-level suppression of lipid metabolism in tauopathy. This pattern is consistent with the broader recognition that disrupted brain cholesterol homeostasis is an early and pathogenically important event in AD, and that miRNA-mediated post-transcriptional suppression of genes may compound the transcriptional downregulation of biosynthetic enzymes identified in our mRNA data.

### Evidence for Brain-Plasma EV Correlation and Its Mechanistic Basis in Tauopathy

A central translational goal of this study was to determine the extent to which plasma EV ncRNA cargo correlates with the central brain transcriptome in tauopathy. Our integration analyses demonstrate that the GO pathway enrichments for pEV miRNA target genes and brain circRNA regulatory networks converge on identical biological themes—specifically genes associated with cholesterol biosynthesis suppression and immune activation in PS19 compared to WT—suggesting a high degree of functional alignment between the brain and peripheral compartments. This brain to plasma EV alignment is mechanistically supported by the findings of Ruan et al.[95, 96], who demonstrated that EVs isolated from human AD brains and injected into wild-type mouse hippocampi could initiate tauopathy, establishing that EVs could be functional carriers of tau-related molecular information. The ncRNA signatures including miR-126a-3p and miR-335-3p which were significantly downregulated in PS19 pEVs compared to WT, further support this link. miR-126-3p is a critical regulator of vascular integrity, and its differential abundance in PS19 compared to WT mice mirrors observations in AD patient plasma derived-EVs where levels correlate with disease severity [3, 67–70, 94]. These cross-compartment correlations provide strong evidence that the ncRNA cargos of circulating plasma EVs effectively track the transcriptional state of the brain in tauopathy.

### Predictive Potential of Plasma EV ncRNA Cargos for AD

The high degree of compartmental convergence and the identification of shared ncRNA species suggest that plasma EV transcriptomics have significant potential for predicting AD pathogenesis and tracking disease progression. miRNAs such as miR-126a-3p and miR-335-3p, which we found significantly downregulated in PS19 pEVs compared to WT, have been validated in human clinical studies as biomarkers that discriminate between AD and healthy controls with high sensitivity and specificity [3, 95–97]. For instance, clinical studies using small RNA sequencing of neuron-derived EVs from human plasma identified miR-126-3p as part of a panel that correlates with temporal cortical thickness on MRI [32, 98] and reflects underlying vascular-to-neuronal signaling deficits. The unique shared signature of brain and pEVS, including miR-5114 and the 32 circRNAs we identified in PS19 compared to WT, provides a specific potential biomarker set that may reflect early central changes prior to overt dementia. Recent evidence suggests that circRNAs act as molecular sponges for microRNAs involved in synaptic plasticity and memory, positioning them not just as markers, but as active participants in the regulation of cognition and cognitive diseases [99]. The identified shared ncRNAs, including miR-5114 and circRNA species like circ_0008242, circ_0002153, circ_0007688 present several advantages as non-invasive early biomarkers. Their presence in EVs ensures protection from systemic RNases, granting them superior stability compared to free-floating transcripts, while their compartmental origin allows them to directly reflect the biochemical status of the CNS in a peripheral biofluid [100–105]. Their compartmental origin—specifically when enriched for brain-specific markers like L1CAM or NCAM or APLP1+ -allows them to directly reflect the biochemical status of the CNS in a peripheral biofluid, bypassing the dilution effects of general plasma RNA [106–112]. Because these molecules participate in the early neuroinflammatory and metabolic transcriptional shifts that preceded neuronal loss, they offer a sensitive diagnostic window for tracking the transition from pre-symptomatic tau accumulation to overt neurodegeneration. In summary, the convergent brain-plasma EV ncRNA signatures identified in this study provide a robust foundation for developing non-invasive molecular tools to monitor the progression of tauopathy.

### Limitations

While this study provides a comprehensive multi-compartment atlas of tauopathy, several limitations must be acknowledged. First, the PS19 model is a model of primary tauopathy and does not capture the complex amyloid-tau synergy present in sporadic human AD. Consequently, the transcriptomic signatures identified here may reflect tau-specific biological processes rather than the full spectrum of AD pathology. Second, this study utilized a single time point (∼9.6 months), capturing a stage of advanced tauopathy. Longitudinal studies across multiple ages would be necessary to determine the temporal dynamics of these ncRNA signatures and identify the earliest detectable peripheral markers of central pathology in PS19 mice compared to WT. Third, our analysis of pEVs focused on the total circulating pool. While EVs are known to cross the blood-brain barrier, the plasma compartment contains a highly heterogeneous mixture of vesicles derived from various tissues, which may dilute CNS-specific signals. Future studies employing immuno-affinity enrichment for CNS-specific markers (e.g., L1CAM) would likely enhance the sensitivity of detecting brain-derived signatures. Finally, while we have identified shared signatures and convergent pathways, functional validation of the identified circRNAs and miR-5114 in primary cell systems is required to elucidate their exact mechanistic roles in tau-mediated neurodegeneration.

## Conclusions

In summary, this study provides the first integrated multi-compartment transcriptomic atlas of the PS19 tauopathy model, demonstrating that cortical tau accumulation triggers a highly coordinated molecular response across brain and peripheral plasma EV compartments. Our analysis revealed a robust transition from homeostatic to DAM gene signature and a systemic suppression of cholesterol biosynthesis transcriptional response in PS19 mice compared to WT. We established that the circular RNA landscape is the most numerically perturbed non-coding RNA biotype in both the cortical tissue and pEVs, identifying a specific signature of 33 shared differentially enriched ncRNAs in PS19 mice compared to WT controls—including miR-5114 and circ_0008242, circ_0002153, circ_0007688 —that consistently reflects central pathology in the periphery. These findings confirm that the non-coding cargo of circulating plasma EVs in PS19 mice effectively reflects central neuroinflammatory and metabolic gene expression profile, providing a robust foundation for the development of non-invasive molecular tools to monitor AD progression and evaluate therapeutic interventions.

## Abbreviations

The following abbreviations are used in this manuscript:

(WT): Wild-Type
(mRNA): messenger Ribonucleic Acid
(ncRNA): non-coding RNA
(pEV): plasma small Extracellular Vesicle
(DEGs): Differentially Expressed Genes
(DAM): Disease-Associated Microglia
(circRNAs): circular RNAs
(miRNA): micro-RNA
(AD): Alzheimer’s Disease
(NFTs): Neurofibrillary Tangles
(Aβ): Amyloid-Beta
(MAPT): Microtubule-Associated Protein Tau
(FTD): Frontotemporal Dementia
(PS19 line): P301S transgenic mouse
(WGCNA): Weighted Gene Co-expression Network Analysis
(snoRNAs): small nucleolar RNAs
(tRNAs): transfer RNAs
(iPSCs): induced Pluripotent Stem Cells
(m6A): N6-methyladenosine
(L1CAM): L1 Cell Adhesion Molecule
(GO): Gene Ontology
(IHC): Immunohistochemistry
(snRNA): small nuclear RNA
(MRPS): Microfluidic Resistive Pulse Sensing
(DAMPs): Damage-Associated Molecular Patterns
(NCAM): Neural Cell Adhesion Molecule
(APLP1+): Amyloid Beta Precursor Like Protein 1
(IACUC): Institutional Animal Care and Use Committee
(ER): Endoplasmic Reticulum
(HRP): Horseradish Peroxidase

## Acknowledgments

We would like to acknowledge Amanda Moore for helping with the pEV and RNA isolation, and library generation.

## Funding

This research was funded by the grants provided to Radosveta Koldamova and Iliya Lefterov, grant number R01 AG077636, R01 AG075992”, and Nicholas Francis Fitz, grant number R01AG075069.

## Author Contributions

Conceptualization, R.K. and I.L.; methodology, T.T.L., A.N.M.M-O-R; software, T.T.L., A.N.M.M-O-R, D.C.L., N.F.F.; validation, T.T.L., A.N.M.M-O-R; formal analysis, T.T.L., A.N.M.M-O-R, D.C.L.; investigation, T.T.L., A.N.M.M-O-R; resources, N.F.F., R.K. and I.L.; data curation, T.T.L., A.N.M.M-O-R; writing—original draft preparation, T.T.L., A.N.M.M-O-R; writing—review and editing, D.C.L., N.F.F., R.K. and I.L.; visualization, T.T.L., A.N.M.M-O-R, D.C.L.; supervision, N.F.F., R.K. and I.L.; project administration, N.F.F., R.K. and I.L.; funding acquisition, N.F.F., R.K. and I.L. All authors have read and agreed to the published version of the manuscript.

## Institutional Review Board Statement

All animal studies were approved by the University of Pittsburgh Institutional Animal Care and Use Committee (Protocol # 24024421, Approved 2/29/2024) and conducted in accordance with the guidelines of the Care and Use of Laboratory Animals.

## Informed Consent Statement

Not applicable.

## Data Availability Statement

Data will be uploaded to GEO upon acceptance. Conflicts of Interest: The authors declare no conflicts of interest.

## Supplementary Materials

N/A.

## Notes

### Competing Interest Statement

The authors have declared no competing interest.

